# Quantifying the effects of anagenetic and cladogenetic evolution

**DOI:** 10.1101/005561

**Authors:** Krzysztof Bartoszek

## Abstract

An ongoing debate in evolutionary biology is whether phenotypic change occurs predominantly around the time of speciation or whether it instead accumulates gradually over time. In this work I propose a general framework incorporating both types of change, quantify the effects of speciational change via the correlation between species and attribute the proportion of change to each type. I discuss results of parameter estimation of Hominoid body size in this light. I derive mathematical formulae related to this problem, the probability generating functions of the number of speciation events along a randomly drawn lineage and from the most recent common ancestor of two randomly chosen tip species for a conditioned Yule tree. Additionally I obtain in closed form the variance of the distance from the root to the most recent common ancestor of two randomly chosen tip species.

## 1 Introduction

One of the major debates in evolutionary biology concerns when evolutionary change takes place. Two predominant theories state that changes take place either at times of speciation [punctuated equilibrium Eldredge and Gould, 1972, Gould and Eldredge, 1993] or gradually over time [phyletic gradualism, see references in Eldredge and Gould, 1972].

Phyletic gradualism describes how Darwin envisioned evolution happening [Eldredge and Gould, 1972]. The theory of punctuated equilibrium was brought up in response to what fossil data was indicating [Eldredge and Gould, 1972, Gould and Eldredge, 1977, 1993]. One rarely observed a complete unbroken fossil series but more often distinct forms separated by long periods of stability [Eldredge and Gould, 1972]. Falconer and Darwin at the birth of the theory of evolution already discussed this and the latter saw “the fossil record more as an embarrassment than as an aid to his theory” [Eldredge and Gould, 1972].

Therefore instead of continuous evolution Eldredge and Gould [1972] developed the idea that change takes place predominantly at speciation in the form of rapid evolution which are effectively jumps at the time scale of most phylogenies [Simpson, 1947]. They were of course not the first to do this. As already mentioned such a theory can be found in Falconer’s writings and was presented under the name “quantum evolution” by Simpson [1947]. In fact Simpson [1947] postulated that this quantum evolution was responsible for the emergence of taxonomic units of high order. However, the work of Eldredge and Gould [1972] re–introduced punctuated equilibrium into contemporary mainstream evolutionary theory.

One of the theoretical motivations behind punctuated equilibrium is that the organism has to function well as a whole. Any particular change to any trait can easily make it dysfunctional with the rest of the organism. Therefore traits have to be adapted to each other first [Chetverikov, 1961, Dobzhansky, 1956, Frazzetta, 1975]. Any initial change to a trait will most probably be inadaptive or non–adaptive first [Simpson, 1947]. One cannot expect the change to be large as then it could make the organism poorly adapted to its environment or the trait would not function well in co–operation with other traits or functions of other body parts. The probability of a mutation with a large positive adaptive phenotypic effect occurring is very small [Simpson, 1947]. However, it might happen that a new environmental niche appears [due to some species dying out and leaving it empty Simpson, 1947] for which the direction of the change is beneficial. Therefore if the organism starts taking up the niche and enhancing this new change, rapid evolution has to take place for all body parts to catch up until a new equilibrium is reached. The alternative [and usual Simpson, 1947] result of such a change is that the organism becomes extinct due to no niche appearing while at the current one the new trait was at a disadvantage or evolution was not rapid enough to fill up the new niche.

Simpson [1947] discussed that it would be easiest for such a change to fixate inside a “small and completely isolated population.” Chetverikov [1961] also postulated this mechanism for differentiation due to isolation inside a species. Mayr [1982] hypothesized that genetic imbalance of isolated subpopulations of a species would favour change and additionally pointed out that this situation is a possible alternative explanation to the observed long periods of stasis in the fossil record. Paleontologists usually encounter widespread populous species and these by the previous argument would be the least likely to change.

Showing whether a trait gradually evolves in a continuous manner or is subject to large rapid changes followed by relative stability is not only a result in itself but gives us actual insights about the role of the trait for the organism. Mooers et al. [1999] discuss this. They expect cladogenetic evolution to occur in traits that are involved in speciation or niche shifts and gradual evolution in traits under continuous selection pressures that often change direction.

Biological evidence for both gradual and punctuated evolution is present. As discussed by Bokma [2002] punctuated equilibrium is supported by fossil records [see Eldredge and Gould, 1972, Gould and Eldredge, 1977, 1993] and as already mentioned they were the motivation for the development of the theory. On the other hand Ozawa [1975]’s study of the Permian verbeekinoid foraminifer *Lepidolina multiseptata* provides evidence for gradual evolution. Stebbins and Ayala [1981] in their work also point to experiments supporting phyletic gradualism. In a more recent work Hunt [2008] found that both a punctuated model and gradual model seemed plausible for evolution of pygidial morphology on a lineage of the trilobite *Flexicalymene.* One should suspect, as Simpson [1947] pointed out, that evolution will be a mix of different modes and interestingly body size evolution in the radiolarian *Pseudocubus vema* and shape components in the foraminifera *Globorotalia* were best explained by a model with both anagenetic and cladogenetic components [Hunt, 2008]. Even more recently Hunt [2013] studied the deep–sea ostracode genus *Poseidonamicus.* Using a maximum likelihood approach he found that only one or two shape traits seem to have evolved via speciational change while four other shape traits evolved gradually (Brownian motion).

Mathematical models with gradual and punctuated components have been considered previously in the literature. Examples of this are due to Bokma [2002, 2008, 2010], Mattila and Bokma [2008], Mooers and Schluter [1998], Mooers et al. [1999]. In a very recent work Eastman et al. [2013] study *Anolis* lizards evolution using a Brownian motion with jumps inside branches model. Testing, especially based on extant species only, whether gradual or punctuated evolution is dominating or if both contribute similarly can be a difficult matter. One only observes the contemporary sample and it is necessary to divide the signal between the two sources of stochasticity. Work has of course been done in this direction. For example Avise [1977] studied the type of evolution in some contemporary fish families and Mattila and Bokma [2008] asked whether gradual or punctuated evolution is responsible for body size evolution in mammals. Hunt [2008] developed a likelihood–based statistical framework for testing whether a purely gradual (Brownian motion) or a purely punctuated model or a mix of these two mechanisms explained phenotypic data best. It should also be remembered that measurement error can make estimation and testing even more difficult. Even with only gradual change, measurement error can have very complicated effects in comparative data e.g. phylogenetic regression coefficients can be both upward or downward biased depending on both the tree and true model parameters [Hansen and Bartoszek, 2012].

The evolutionary model considered in this work has two main components. One is the model of phenotypic evolution and the other the branching process underlying the phylogeny. Currently most of the phylogenetic comparative literature is set in a framework where one conditions on a known tree. Whilst with the current wealth of genetic data which allows for more and more accurate phylogenies this is a logical framework, one can run into situations where this is not sufficient. Typical examples are fossil data or unresolved clades e.g. in the Carnivora order [used for an example analysis in Crawford and Suchard, 2013]. To add to this, very recently a new member of the olingos (*Bassaricyon*, Order Carnivora) the Olinguito (*B. neblina*) has been described [Helgen et al., 2013]. Moreover when considering models with a jump component one has to consider speciation events leading to extinct lineages and how this interplays with the model of phenotypic evolution assumed here. The tree and speciation events themselves might not be of direct interest, in fact they could actually be nuisance parameters. So there is interest in tree–free methods that preserve distributional properties of the observed phenotypic values [Bokma, 2010].

I consider in this work two models of phenotypic evolution. Brownian motion, interpreted as unconstrained neutral evolution and a single optimum Ornstein–Uhlenbeck model. In the case of the second model the phenotype tends to constrained oscillations around the attractor. On top of this just after a speciation event each daughter lineage undergoes rapid change, described here by some particular probability density function.

Combining the Ornstein–Uhlenbeck process with a jump component makes phylogenetic comparative models consistent with the original motivation behind punctuated equilibrium. After a change occurs rapid adaptation of an organism to the new situation takes place followed by stasis. Stasis as underlined by Gould and Eldredge [1993] does not mean that no change occurs. Rather that during it “fluctuations of little or no accumulated consequence” occur. The Ornstein–Uhlenbeck process fits into this idea. If the adaptation rate is large enough then the process reaches stationarity very quickly and oscillates around the optimal state and this can be interpreted as stasis between the jumps — the small fluctuations. Mayr [1982] supports this sort of reasoning by hypothesizing that “The further removed in time a species from the original speciation event that originated it, the more its genotype will have become stabilized and the more it is likely to resist change.”

Literature concerning punctuated equilibrium models is predominantly concentrated on applications and estimation. Here I take a more theoretical approach and study the mathematical properties of evolutionary models with an additional component of rapid phenotypic change associated with speciation assuming that all model parameters are known (or have been preestimated). To the best of my knowledge an adaptive evolutionary model with a punctuated component has not been developed fully. The main contribution of this paper is the mathematical development of such a model — the Ornstein–Uhlenbeck model with jumps.

Current Ornstein–Uhlenbeck based evolutionary models have a discontinuity component but this is in the primary optimum [Bartoszek et al., 2012, Butler and King, 2004, Hansen, 1997]. The phenotype evolves gradually towards oscillations around a new optimum value and this can correspond to a change in the environment, a sudden replacement of the adaptive niche. The model considered here is different. It is the phenotype which jumps, it is pushed away from its optimum due to e.g. a mutation that changed the character dramatically. After the jump the phenotype evolves back towards the optimum.

Of course it would be more realistic to have a discontinuity in both the phenotype and the primary optimum. Then when the jump in the phenotype and optimum function would coincide the species would be at an advantage needing a shorter time to reach equilibrium. However, this setup would require some sort of model for the optimum and is beyond the scope of this study.

The other model considered by us is a Brownian motion model with jumps. It differs from the Ornstein–Uhlenbeck one in that it does not have an adaptive constraint. In fact, as I show in my mathematical derivations, it is the limit as the rate of adaptation goes to 0. From a biological point of view this model illustrates unconstrained evolution with rapid speciational change. Additionally it serves as a warm–up to the mathematics of the significantly more involved Ornstein–Uhlenbeck model.

The Ornstein–Uhlenbeck with jumps model combines three important evolutionary components. Firstly it has built into it a phylogenetic inertia component, the ancestral value of the trait will make up part of the contemporary species’s trait value and additionally cause dependencies between current species. Secondly unlike the Brownian motion model the variance of this process tends to a constant and so after some time the trait should exhibit constrained oscillations around the optimum value – stasis. And finally there is the jump component — the possibility that at speciation the trait undergoes rapid evolution.

In this work apart from introducing a model combining adaptation and punctuated equilibrium I propose a way of quantitatively assessing the effect of both types of evolution so that one can consider a mix of gradual and punctuated components. The analytical results are computed for a pure birth model of tree growth, however using example data I show that they can be carried over (at the moment via simulation methods) to branching process models that include extinction.

The phylogeny, number and timing of speciation events, is modelled by a constant–rates birth–death process conditioned, on the number of tip species, *n*. By λ I denote the birth rate and by *μ* the death rate. Contemporary literature on this is substantial [e.g. Aldous and Popovic, 2005, Gernhard, 2008a, Mooers et al., 2012, Stadler, 2009, 2011, Stadler and Steel, 2012]. The key mathematical property of conditioned branching processes is that conditional on the tree’s height the times of speciation events are independent and identically distributed random variables. The distribution is particular to the regime: critical (λ = *μ*), supercritical (λ > *μ* > 0) or pure birth (λ > *μ* = 0). Estimation of birth and death rates has been widely discussed in the literature [e.g. Bokma, 2002, 2003, Bokma et al., 2012, Nee, 2001, Nee et al., 1994a, b].

The combination of evolutionary processes and conditioned branching processes has already been considered by Edwards [1970] who proposed a joint maximum likelihood estimation procedure of a pure birth tree and a Brownian motion on top of it. Markov–Chain Monte Carlo based methods to jointly estimate the phylogeny and parameters of a Brownian motion trait have been proposed by Huelsenbeck et al. [2000] and Huelsenbeck and Rannala [2003]. Slater et al. [2012] develop an Approximate Bayesian Computation framework to estimate Brownian motion parameters in the case of an incomplete tree. Sagitov and Bartoszek [2012], Crawford and Suchard [2013], Bartoszek and Sagitov [2012] have contributed by considering a Brownian motion on an unobserved birth–death tree in the first two works and an Ornstein–Uhlenbeck process on an unobserved pure birth tree in the third. These studies concentrate on the speciation process driving the phenotypic process, as is usually assumed in phylogenetic comparative methods, where the trait is of main interest. There have been a number of recent papers concerning models where the rate of speciation depends on the trait values. Pie and Weitz [2005] discuss many possible modelling examples where the phenotypic and speciation processes interact. Maddison et al. [2007] derive likelihood formulae for a model where a binary character drives speciation and extinction and study this model via simulations. Closer to the present setting is FitzJohn [2010] who assumed the trait evolved as a diffusive process with birth and death coefficients as functions of the trait. A similar framework of combining Ornstein–Uhlenbeck process and branching processes has been considered by Rossbert et al. [2010] and by Brännstrröm et al. [2011] in the food-web structure community.

The organization of this paper is as following. In Section 2 I discuss the pure birth model of tree growth, derive the probability generating function of the number of speciation events on a random lineage and from the most recent common ancestor of two randomly sampled tip species and in addition also show how my results can be applied to working with the total tree area. Section 3 is devoted to punctuated equilibrium evolutionary models. Section 4 interprets the results of the Hominoidea analyses of Bokma [2002], Bokma et al. [2012] in the light of the present models and Section 5 is a discussion, Appendix A contains proofs of the main mathematical results Theorems 2.1 and 3.2, and Appendix B presents a brief introduction to quadratic variation.

## 2 The conditioned Yule process

### 2.1 Yule model of tree growth

The Yule tree model [Yule, 1924] is a pure birth Markov branching process. At the beginning there is one species that lives for an exponential time and then splits into two species each behaving in the same manner. Here I will consider a Yule process conditioned on *n* tips: the extant species These types of branching processes, called conditioned birth–death processes, have received significant attention in the last decade [e.g. Aldous and Popovic, 2005, Gernhard, 2008a, Mooers et al., 2012, Stadler, 2009, 2011, Stadler and Steel, 2012].

For the purpose of this current work I need the Laplace transform of, *T*, the height of a Yule tree conditioned on *n* tips at present and the Laplace transform of the random variable *τ*, the time to the most recent common ancestor of two species randomly sampled out of *n*. These have already been derived and studied in detail by Bartoszek and Sagitov [2012]. In addition to be able to incorporate the jump events I will need to study the random variables ϒ and *ν* — the number of speciation events from the time of origin of the tree until a randomly chosen tip species and the number of speciation events that occurred on the lineage from the tree origin to the most recent common ancestor (excluding it) of two randomly sampled tip species respectively. Figure 1 visualizes these random variables. These two random variables can also be seen as distances on the tree counted as number of edges, ϒ — distance from the root of random tip, *ν* — distance from the root of the most recent common ancestor of two randomly sampled tips.

**Figure 1.**
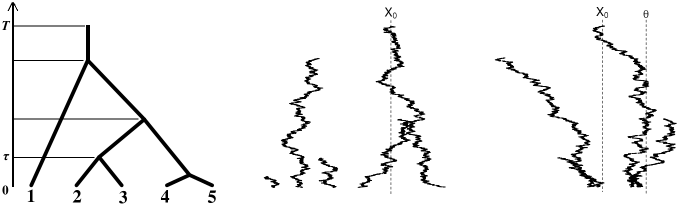
Left: a conditioned Yule tree (λ =1, *n* = 5) with the different time components marked on it. The height of the tree is *T*. If species 2 is “randomly” chosen then it would have ϒ = 3 and if the pair of species 2, 3 is “randomly” chosen then they would have *ν* = 2 and the time till their coalescent would be *τ*. Center: a Brownian motion with jumps 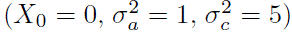 evolving on the tree and right: an Ornstein–Uhlenbeck process with jumps (*α* = 1, *θ* = 1, 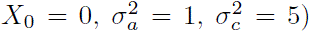 evolving on the tree. Simulations were done using the TreeSim [Stadler, 2009, 2011] and mvSLOUCH [Bartoszek et al., 2012] R [R Core Team, 2013] packages.

**Figure 2.**
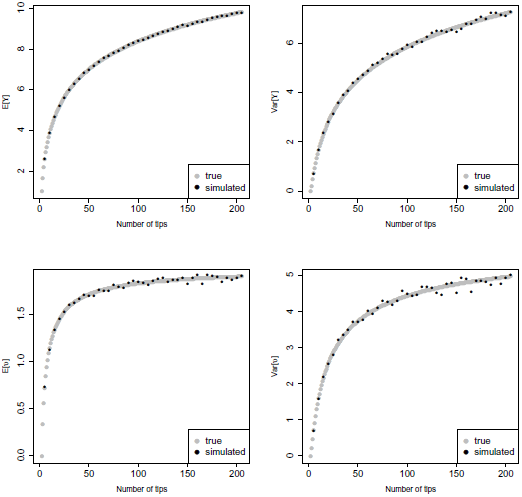
Simulated and true values of E[ϒ], Var[ϒ] (top) and E[*ν*], Var[*ν*] (bottom). Each point comes from 10000 simulated Yule trees. Var[*ν*] grows very slowly and with *n* = 200 it is still rather distant from its asymptotic value of 6.

**Figure 3.**
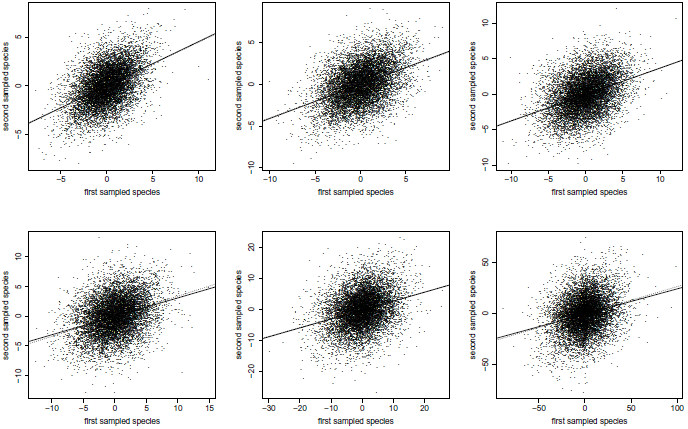
Regression lines fitted to simulated data (thick line) mostly indistinguishable from the true regression line *y* = *ρ*_*n*_*x* (dotted line) with *ρ*_*n*_ given by the exact formula (Eq. 18) for different values of *κ* in the Yule–Brownian–motion–jumps model. Top row from left to right: κ = 0.0099,0.3333, 0.5, bottom row from left to right: *κ* = 0.6667, 0.9091,0.9901. In all cases the jump is normally distributed with mean 0 and variance 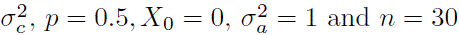.

**Figure 4.**
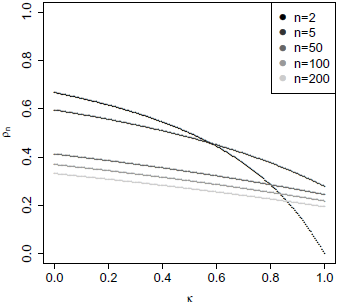
Interspecies correlation coefficient for the Yule–Brownian–motion–jumps model for different values of *n*.

**Figure 5.**
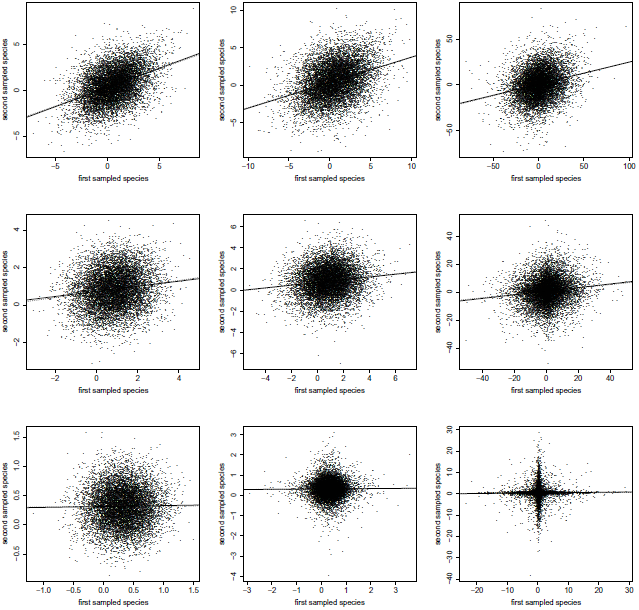
Regression lines fitted to simulated data (thick line) mostly indistinguishable from the true regression line 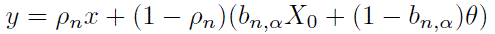 (dotted line) with *ρ*_*n*_ given by the exact formula for different values of *κ*, *α* and *δ* in the Yule–Ornstein–Uhlenbeck–jumps model. Top row *α* = 0.05, center row *α* = 0.5, bottom row *α* = 5. First column *κ* = 0.01 second column *κ* = 0.5 and third column *κ* = 0.99. The other parameters are fixed at 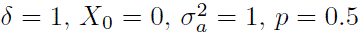, so 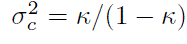.

I recall the following key lemma concerning a Yule tree,

#### Lemma 2.1

(Bartoszek and Sagitov, 2012, Stadler, 2009). *In a Yule tree conditioned on n contemporary species the probability that the coalescent of two randomly sampled tips occurred at the *k*–th (counting forward in time) speciation event is,*

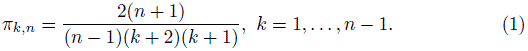

I use the following notation,

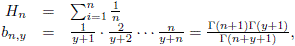

where Γ(⋅) is the gamma function. In the special case *y* = *j*, where *j* is an integer, 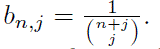 Using this I recall [from Bartoszek and Sagitov, 2012] the
Laplace transforms of T and *τ*,

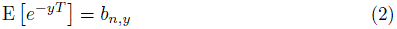

and

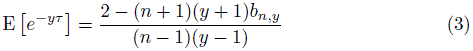

with 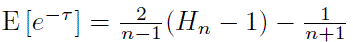 as the limit of *y* → 1.

I further recall [Bartoszek and Sagitov, 2012, Sagitov and Bartoszek, 2012, Steel and McKenzie, 2001],

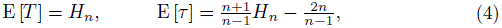

both behaving as ln *n* [however see also Gernhard, 2008a, 2008b, Mooers et al., 2012, Sagitov and Bartoszek, 2012, Stadler, 2008, Stadler and Steel, 2012, for situations with extinction present].

### 2.2 Counting speciation events

I defined the random variable ϒ as the number of speciation events from the time of origin of the tree until a randomly chosen tip species (see Fig. 1). I can write 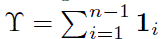, where **1**_*i*_ is the indicator random variable of whether the *i*-th speciation event (counting from the first speciation event) is on the randomly chosen lineage. The probability that **1**_*i*_ = 1 is 2/(*i* + 1). The argument behind this is as follows, just before the *i*-th speciation event there are *i* + 1 points that need to coalesce. Exactly one of these is on the lineage of interest. There are 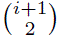 in total possibilities of choosing the two points that will coalesce. Exactly *i* will contain the point of interest, so as I consider the Yule tree model the probability is 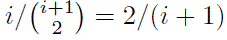. From this I get that

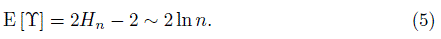

Now let us consider the situation that I sample two tip species and am interested in the expectation of, *ν*, the number of speciation events that occurred on the lineage from the tree origin to their most recent common ancestor (excluding it), see Fig. 1. Using Eq. (1) and the following identity,

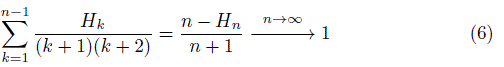

this equals,

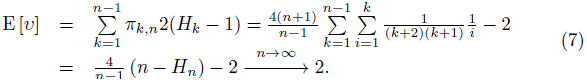

The same formulae for E[ϒ] and E[*ν*] were derived by Steel and McKenzie [2001] alongside the distribution and variance of ϒ.

I can add a variation to the above discussion by marking each speciation event with probability *p*. Let then ϒ* ≤ ϒ count the number of marked events along a randomly chosen lineage. Obviously

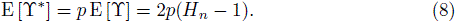

Similarly let *ν*\* ≤ *ν* count the number of marked speciation events from the tree origin to the most recent common ancestor of two randomly chosen tip species. Again

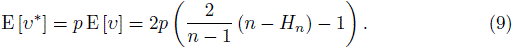

I further derive here the probability generating functions of ϒ and *ν*. They do not depend on the speciation rate λ (a scale parameter for branch lengths when conditioning on *n*) as ϒ
 and *ν* are topological properties. Notice in the theorem below that *b*_*n*-1,2*s*-1_ is well defined because 2*s* − 1 > −1 (as 2*s* > 0) and therefore consequently the probability generating functions are well defined for all *s* > 0. One should also not forget that in the case of there being only two extant species *ν* equals 0 due to there being only one speciation event — the most recent common ancestor of the only pair of species — and this is not counted by *ν*.

#### Theorem 2.1.

*The probability generating function for* ϒ *is*,

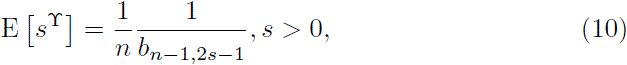

*while for *ν**,

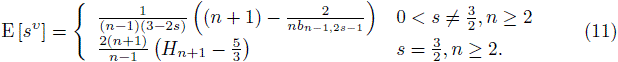

Using the value of the second derivatives of the probability generating function at 1 I can calculate the variance of ϒ [obtained earlier in Steel and McKenzie, 2001, in a different manner] and u as,

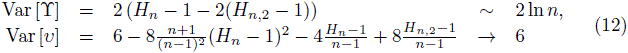

where 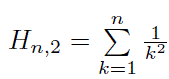.

#### 2.2.1 Connection to total tree area

The *total area* of a tree *T* with *n* tips, is defined [as in Mir and Rosselló, 2010],

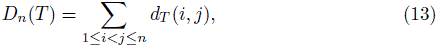

where *d_T_*(*i*,*j*) is the distance between nodes *i* and *j*. This distance can be counted in two ways, either as the number of edges on the path between the two nodes or as the number of vertices. The latter will be one less than the former. For a Yule tree I can calculate the expectation of the total area as

- 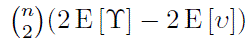 with the first definition of *d_T_*(*i*, *j*),
- 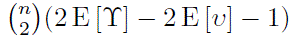 with the second definition of *d_T_*(*i*, *j*).

Plugging in the values for E[ϒ] and E[*ν*], Eqs. (5) and (7), I get,

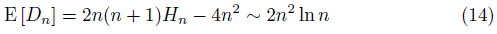

and

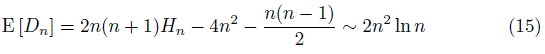

respectively, depending on the definition of *d_T_*(*i*, *j*). These results are the same as in the literature [Mir et al., 2013, in the case of the first definition] and [Mulder, 2011, in the case of the second definition]. Mir et al. [2013] claim that the expectation of the total area of a Yule tree calculated by Mulder [2011] contains an error, however the discrepancy between the results of these two studies comes from them using the different definitions of the distance between two tips.

## 3 Models with punctuated evolution

Stochastic models for continuous trait evolution are commonly based on a stochastic differential equation (SDE) of the type,

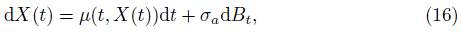

along the phylogenetic tree [see e.g. Bartoszek et al., 2012, Butler and King, 2004, Felsenstein, 1985, Hansen, 1997, Hansen et al., 2008, Labra et al., 2009] At speciation times this process divides into two processes evolving independently from that point. Very often it is assumed that the natural logarithm of the trait evolves according to this SDE [for some motivation for this see e.g. Bartoszek et al., 2012, Bokma, 2002, Huxley, 1932, Savageau, 1979]

It is straightforward to include in this framework a mechanism for modelling cladogenetic evolution. I consider two possible mechanisms. The first one is that just after each speciation point in each daughter lineage with probability *p* a jump (mean 0, variance 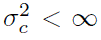) takes place. The second mechanism is that at each speciation point one adds to the phenotype process of a randomly (with probability 0.5) chosen daughter lineage a mean 0, variance 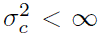 jump. The other daughter lineage is then not affected by the jump. This can be interpreted as some change in the newborn species that drove the species apart. At the present second–moment level of analysis this model is equivalent to the first one with *p* = 0.5. One cannot distinguish between these two punctuated change mechanisms unless one actually observes the speciation and jump patterns. A jump at each daughter lineage with probability 0.5 and a jump in exactly one randomly chosen daughter lineage will have the same effect on the first and second sample moments. This is because they only depend on the expectation and variance of the jump and number of jumps in the common part of the two lineages. In the simulations shown in this work I assume normality of the jump but all the results will hold for any mean 0, finite variance jump.

I will study two currently standard evolutionary models the Brownian motion and Ornstein–Uhlenbeck process both expanded to include a punctuated equilibrium component. In line with previous work [Bartoszek and Sagitov, 2012, Sagitov and Bartoszek, 2012] I do not condition on a given phylogenetic tree but assume a branching process (here conditioned Yule tree) for the phylogenetic tree. In such a case, a number of relevant model properties [see Bartoszek and Sagitov, 2012, Sagitov and Bartoszek, 2012] can be conveniently described in terms of the variance of the trait of a randomly sampled tip species and the covariance (or correlation) between two randomly sampled tip species. This correlation [not to be confused with correlation between traits which can result from developmental constraints Cheverud, 1984] called the interspecies correlation coefficient [Sagitov and Bartoszek, 2012] is a consequence of phylogenetic inertia. Extant species will be correlated due to their shared ancestry. How strong that correlation is will depend on how much time has passed since their divergence (stochastic tree model) and mode of evolution (stochastic trait model).

The interspecies correlation is similar to what Cheverud et al. [1985] describe as the phylogenetic autocorrelation. Cheverud et al. [1985] repeated after Riedl [1978] and Waddington [1957] that the similarity of a trait between species can be indicative of its functional role. A highly central trait should be significantly correlated while a peripheral one should have more freedom to vary. This is in line with thinking of all of the traits interacting with each other. If a trait is central then all the other traits will have adapted to working with it and any change in it would cause all the other traits to be away from their optimum and hence potentially cause a significantly larger misadaptation than change in a trait that is not so central.

Below I discuss the correlation function (with detailed derivations in Appendix A). It will turn out that in the case of punctuated evolution a relevant parameter is 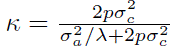. Knowing whether the gradual or jump component dominates tells us whether the correlation is due to shared branch length (gradual dominates) or shared number of speciation events (jumps dominate). One can also recognize a similarity to measurement error theory. If one thinks of the jumps as “errors” added to the trajectory of the evolving diffusion process then 1 − *κ* could be thought of as the reliability ratio or measurement error correction factor [see e.g. Buonaccorsi, 2010, Fuller, 1987, Hansen and Bartoszek, 2012]

The work here is also another step in introducing Lévy processes to the field of phylogenetic comparative methods [see also Bartoszek, 2012, Landis et al., 2013] The framework of Lévy processes is very appealing as it naturally includes punctuated change, i.e. jumps in the evolution of continuous traits.

I introduce the following notation, by *X(t)* I will mean the phenotype process, *X* will denote the trait value of randomly sampled tip species, while *X*_1_ and *X*_2_ will denote the trait values of a randomly sampled pair of tip species.

### 3.1 Unconstrained evolutionary model

The Brownian motion model [Felsenstein, 1985] can be described by the following SDE,

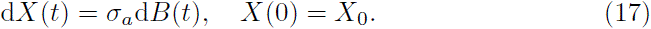

On top of this at each speciation point, each daughter lineage has with probability *p* a mean 0, variance 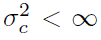 jump added to it. As the jump is mean zero it does not change the expectation of the value of a tip species. Below I consider the variance of a randomly sampled tip species and the covariance and correlation between two randomly sampled tip species.

To derive the following theorem I rely on previous results [Bartoszek and Sagitov, 2012, Sagitov and Bartoszek, 2012]

#### Theorem 3.1.

*The interspecies correlation coefficient for a phenotype evolving as a Brownian motion with jumps on top of a conditioned Yule tree with speciation rate λ =* 1 *is,*

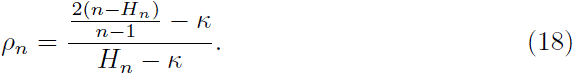

#### Proof.

Due to the jumps being independent of the evolving Brownian motion process

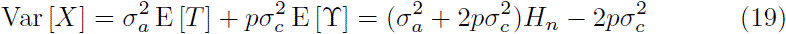

and the covariance between two randomly sampled tip species,

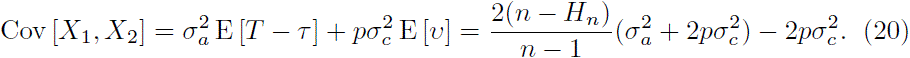

Taking the quotient of these two values results in the desired formula for *ρ*_*n*_.

Comparing with the correlation coefficient for the Brownian motion process calculated by Sagitov and Bartoszek [2012] adding jumps causes both the numerator and denominator to be corrected by *κ*. The following asymptotic behaviour can be directly seen,

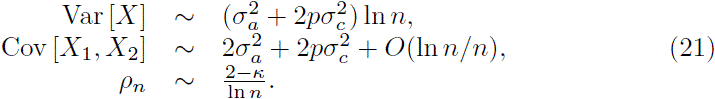

Taking the derivative of the correlation in terms of *κ* I find that it is negative and so:

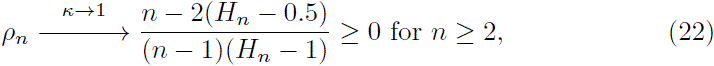

which is a monotonically decreasing convergence.

In the above I assumed that the speciation rate is λ = 1. This restriction does not change the validity of the results as changing λ is equivalent to rescaling the branch lengths by its inverse. As mentioned because I have conditioned on *n* this has no effect on the topology, and therefore does not effect ϒ and *ν*. Consequently a Yule–Brownian–motion–jumps model for the extant species trait sample with parameters 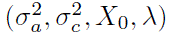 is equivalent to one with parameters 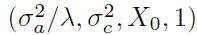.

### 3.2 Constrained model

The basic stochastic process used to model adaptation in the phylogenetic comparative methods field is the Ornstein–Uhlenbeck process [Butler and King, 2004, Felsenstein, 1988, Hansen, 1997, Hansen et al., 2008, Labra et al., 2009],

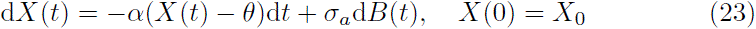

and to it I introduce the jump component in the same fashion. At each speciation point a randomly chosen daughter lineage has a mean 0, variance 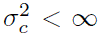 jump added to it. As discussed in the introduction this model is appealing as it allows one to combine gradual and punctuated evolution in one framework.

However, in this model a jump could also affect the mean value and so a more careful treatment is necessary, see the proof in Appendix A. Again I can assume λ = 1 as the following two parameter sets are equivalent, 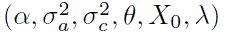 and 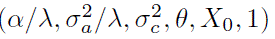 for a Yule–Ornstein–Uhlenbeck–jump model.

To describe the Yule–Ornstein–Uhlenbeck–jump model I introduce the following convenient parameter, 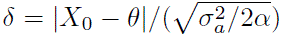 (distance of the starting value from the optimal one scaled by the stationary standard deviation of the Ornstein–Uhlenbeck process).

#### Theorem 3.2.

*The mean, variance, covariance, correlation values in a Yule–Ornstein–Uhlenbeck–jump model with λ =* 1 *are,*

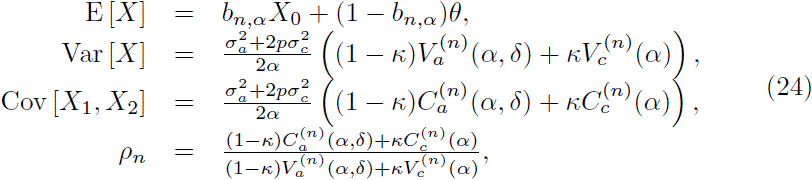

*where,*

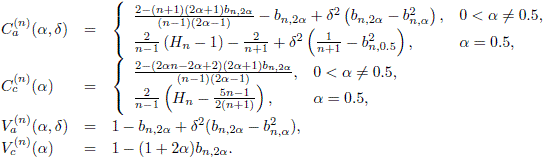

The exact final formula in Theorem 3.2 depends on whether *α* = 0.5 or *α*≠0.5 [see Bartoszek and Sagitov, 2012, for a discussion on this]. Var [*X*] and Cov [*X*_1_, *X*_2_] are made up of two distinct components: one from the Ornstein–Uhlenbeck anagenetic evolution and the other from the cladogenetic “jump” evolution.

Using that as 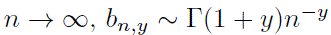 due to the behaviour of the beta function 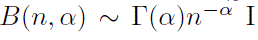 obtain the following asymptotic behaviour for the variance,

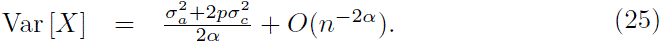

As *α* → 0 by expanding the *b*_*n,y*_ symbol I arrive at the limit,

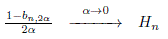

and by this the variance converges on that of a Yule–Brownian–motion–jumps model as *α* → 0. Additionally using the de L’Hôspital rule I obtain that the covariance converges on that of a Yule–Brownian–motion–jumps model as *α* → 0 and as *n* → ∞ the covariance behaves as,

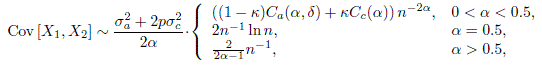

where

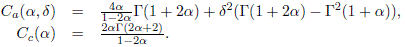

Asymptotically as *n* → ∞ the correlation coefficient behaves (depending on *α*) as,

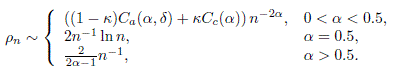

Depending on whether 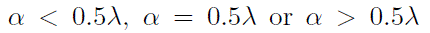 (remember the model equivalency with λ ≠ 1) there are different asymptotic regimes. This has been also noticed by Adamczak and Miłoś [2011, in press] and Bartoszek and Sagitov [2012]. An intuitive explanation why for *α* < 0.5 a completely different behaviour occurs can be that in this case the branching rate is relatively high (with respect to *α*) and local correlations will dominate over the ergodic properties of the Ornstein–Uhlenbeck process [Adamczak and Miłoś, 2011, in press]. However, why this threshold lies at exactly *α* = 0.5λ remains unclear.

As *κ* → 1 the correlation coefficient converges to 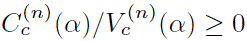. With fixed *α* and *n* one can immediately see that this has to be a monotonic, either increasing or decreasing convergence. Because a jump component adds independent of the trait value noise to the system one can expect it to be a decreasing convergence, and plotting the correlation for different values of the remaining parameters confirms this, (Fig. 6). However, a full mathematical proof is still lacking due to the delicate interactions of the different components of *ρ*_*n*_(*κ*). The conjecture stated below gives us the equivalent condition for the interspecies correlation coefficient *ρ*_*n*_(*κ*) to decrease monotonically for *κ* ∈ (0,1). It is enough to consider *δ* = 0, as for all *n* ≥ 2 and *α*, *δ* ≥ 0 I have, 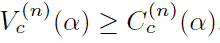.

**Figure 6.**
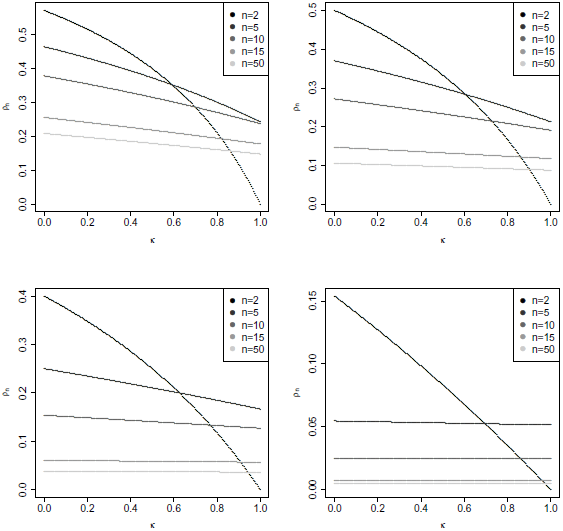
Interspecies correlation coefficient for the Yule–Ornstein–Uhlenbeck–jumps model for *δ* = 0 and different values of *α*, *n*. Top left: *α* = 0.25, top right: *α* = 0.5, bottom left: *α* = 1, bottom right *α* = 5.

**Figure 7.**
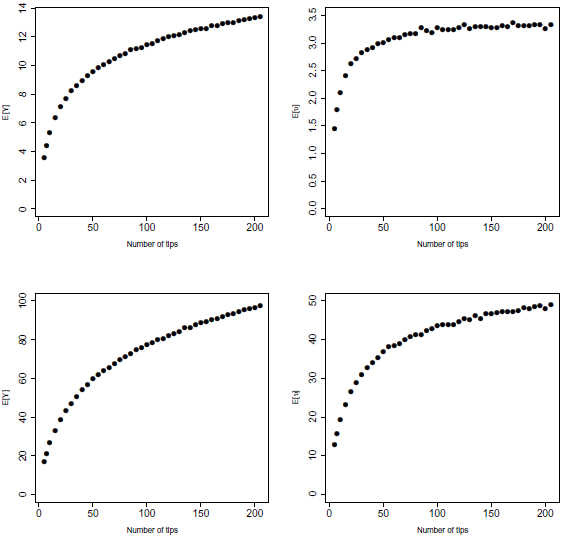
Simulated E[ϒ] and E[*ν*] as functions of *n*, the number of extant species, in the case of a supercritical birth–death tree with λ = 0.134 and *μ* = 0.037 top row (10000 simulated trees) and λ = 0.46, *μ* = 0.43 bottom row (10000 simulated trees). Simulations were done using the TreeSim [Stadler, 2009, 2011] R package.

#### Conjecture 3.1.

*For all α* ≥ 0, *n* ≥ 2

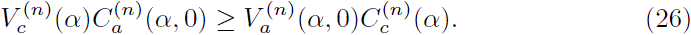

### 3.3 Introducing extinction

Above I concentrated on the case of pure birth trees. However, a more general version of the derived formulae can be used to include death events. For the unconstrained evolutionary model i.e. Brownian motion, I found that:

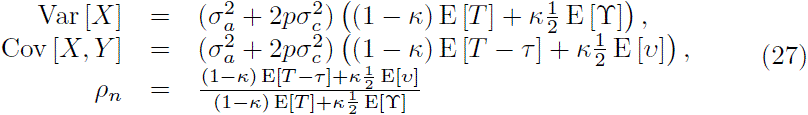

and for the constrained model, with Ornstein–Uhlenbeck dynamics:

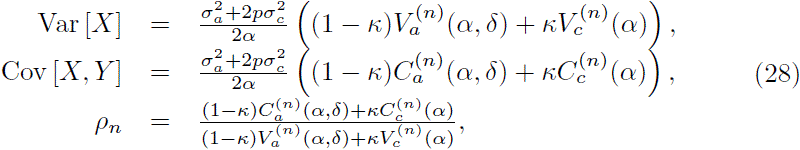

where *p* is the probability of a jump occurring and,

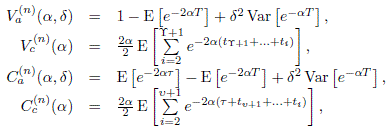

where (*t*_1_,…, *t*_ϒ+1_) are the times between speciation events on a randomly chosen lineage, see Fig. 8.

**Figure 8.**
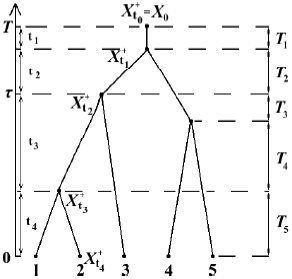
A pure–birth tree with the various time components marked on it. If I “randomly sample” node two then I will have ϒ = 3 and 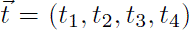. I have 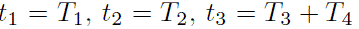 and *t*_4_ = *T*_5_. The process values just after the nodes are 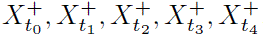, with 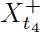 equalling the extant value *X*_2_ if node 2 did not split or if node 2 split 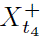 would be *X*_2_ plus a jump (if one took place). If I “randomly sample” the pair of extant species 2 and 3 then *ν* = 1 and the two nodes coalesced when the process was just before 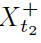.

The values of E[*T*] and E[*T* − *τ*] for a birth–death tree with constant coefficients λ ≥ *μ* > 0 where discussed by Sagitov and Bartoszek [2012] [see also Gernhard, 2008a, b, Mooers et al., 2012, Stadler, 2008, Stadler and Steel, 2012] The formulae for the Laplace transforms of *T* and *τ* and also for the expectation and probability generating functions of ϒ and *ν* are to the best of my knowledge not available yet [but see Mooers et al., 2012, Stadler and Steel, 2012, for some distributional properties], however they could be obtained via simulation methods [using e.g. the TreeSim R package Stadler, 2009, 2011] When discussing the Hominoid body–size analysis of Bokma [2002] below I will illustrate this approach.

### 3.4 Quantifying effects of gradual and punctuated change — quadratic variation

In the previous section I described first– and second–order properties (mean, variance, covariance, correlation) of an evolutionary model containing both gradual and punctuated change. One may ask what is the main effect of cladogenetic evolution on the phylogenetic sample and also how one can elegantly summarize the magnitude of its effect. From the sample’s point of view one can see (if Conjecture 3.1 is correct) that increasing the variance of the jump with respect to the stochastic perturbation of the phenotype will decorrelate the contemporary observations, i.e. they will be less similar than expected from gradual change. At first glance this might seem like a “nearly” obvious statement but notice that the covariance is also increased due to the jump component and all jumps, apart from those on pendant branches, are shared by some subclade of species. Therefore a large jump early enough (to be shared by a large enough subclade) could “give similarity” to these tips.

The decrease of similarity between species with *κ* (conditional on Conjecture 3.1) is the main effect of cladogenetic change but it would be biologically useful to have some value characterizing the magnitude of its effect. One such possibility is the expectation of the quadratic variation (see Appendix B for a brief mathematical introduction to this) of the evolutionary process along a single lineage. As one can attach mechanistic interpretations to the different modes of evolution [Mooers et al., 1999] appropriately partitioning the reasons for evolution could give insights into the trait’s role.

The two models of anagenetic evolution considered here have the same quadratic variation 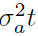. The cladogenetic component comes in as a mean 0, variance 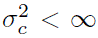 jump just after a speciation event added with probability *p*. Therefore for both of these models (and in fact for any one defined by Eq. 16) with the additional jump component the quadratic variation of a given lineage will be

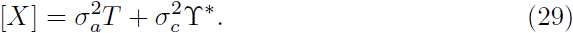

One can clearly see that the magnitude of the trait’s fluctuations can be divided into the cladogenetic and anagenetic component and they are defined respectively by the parameters 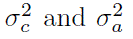 [as considered in Bokma, 2002]. However, the formula for the quadratic variation depends on *T* and ϒ*. These are random unless the tree is given and the jump pattern known. Therefore this is a random variable and instead I consider its expectation,

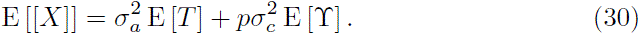

With a given phylogeny one might consider that using E[*T*] is superfluous but this could be useful if one wants to make predictive statements not dependent on the given phylogeny. Under the considered conditioned Yule tree I will have,

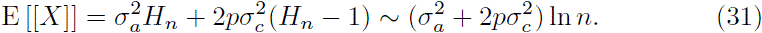

One way of quantifying the proportion of cladogenetic change is using the value

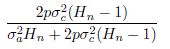

(and respectively

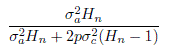

for the anagenetic effects). For a large enough set of tip species the above would simplify to *κ* (and 1 − *κ*).

Mattila and Bokma [2008] proposed two other ways of comparing anagenetic and cladogenetic effects for Brownian motion punctuated evolution. The first one is less relevant here as it involves an estimation procedure. One sets 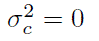 0 and estimates 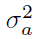 under this assumption. Next one estimates both parameters and compares by how much has the estimate of 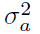 dropped. The second is very similar to the proposed quadratic variation approach. Knowing the speciation and extinction rates the average species lifetime or average time to the next split is known [Bokma et al., 2012, Mooers et al., 2012, Stadler and Steel, 2012, Steel and Mooers, 2010] In the conditioned Yule tree framework used here the length, *t*_*b*_, of a random interior edge is exponential with rate 2λ and hence will have expected length 1/2λ [Mooers et al., 2012, Stadler and Steel, 2012, Steel and Mooers, 2010] Therefore between speciation points the percentage of evolution due to the gradual component will be,

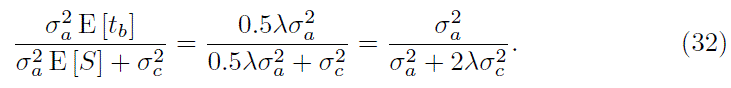

Notice that this will correspond exactly to the 1 − *κ* limit derived in the previous paragraph after one small modification. Namely I assume that the tree starts at the moment of the first speciation event, i.e. disregarding the root branch. Then E[*T*] = *H_n_* − 1 and for all *n* the proportion of anagenetic change will be 1 − *κ* (and cladogenetic *κ*). Taking *p* = 1 i.e. a jump occurred at each speciation event I arrive at Eq. (32).

These two procedures were described by Mattila and Bokma [2008] for a Brownian motion model of gradual evolution. However, the discussion concerning quadratic variation above suggests that they are valid for any model described by Eq. (16).

## 4 Interpreting Hominoid body size evolution results

For illustrative purpose I will now discuss how the conclusions of Bokma [2002] look under my approach. I realize that this clade is not the most appropriate one to be analyzed by a tree–free model because its phylogeny is available. I nevertheless chose to use it here for a number of reasons. First birth and death rate parameters and also the rates of anagenetic and cladogenetic evolution have already been preestimated. I mentioned in the introduction that I do not consider estimation of parameters in this current work but want to concentrate on understanding their effects and using a clade with a known phylogeny makes this easier. One can also use my results if only the time to the most recent common ancestor of the clade is known and I discuss such a situation here.

I do not claim that what I write here about Hominoid body size evolution is authoritative but rather wish to motivate the usefulness and applicability of the methods presented above. In the subsequent discussion I assume that the unit of time is 1 million years.

Bokma [2002] studied the evolution of (the logarithm of) Hominoid body size under a Brownian motion model of gradual evolution (diffusion coefficient 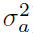) with an independent normally distributed mean 0, variance 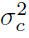 jump added to a newborn daughter lineage at the speciation instant. This is equivalent to the presented here model with *p* = 0.5 [“We can only state in retrospect that the probability that a speciation event has affected the phenotype of interest is 1/2.” Bokma, 2002]

Bokma [2002, see Fig. 2 therein] considers the Hominoidea phylogeny of Purvis [1995] consisting of five species, *Gorilla gorilla, Homo sapiens, Pongo Pygmaeus, Pan paniscus* and *Pan troglodytes.* Purvis et al. [1995] estimated a birth rate of λ = 0.134 and death rate *μ* = 0.037. Other rates could be more appropriate as Bokma et al. [2012] obtained λ = 0.46 (95% CI 0.12 − 1.37) and *μ* = 0.43 (95% CI 0.12 − 1.37) for a seven species phylogeny (*Pan paniscus, Pan troglodytes, Homo sapiens, Gorilla gorilla, Gorilla beringei, Pongo abelii, Pongo pygmaeus*).

Using the phylogeny of Purvis [1995], Bokma [2002] obtained parameter estimates, 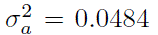 and 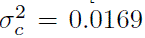. However, a 95% confidence interval for 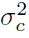 contained 0 but this, as Bokma [2002] pointed out, could be caused by a lack of statistical power due to the low number of extant species.

I derived E[ϒ] and E[*ν*] only for the pure birth process and so Eq. (18) is known only in such a situation. However, in this specific analysis with known (preestimated) λ, *μ* and fixed number of extant nodes I can obtain estimates of E[ϒ] and E[*ν*] via simulations using the TreeSim [Stadler, 2009, 2011] R package. In Fig. 7 one can see the estimates of E[ϒ] and E[*ν*] for different values of *n*. I calculate the values of E[*T*] and E [*T* − *τ*] according to Sagitov and Bartoszek [2012]:

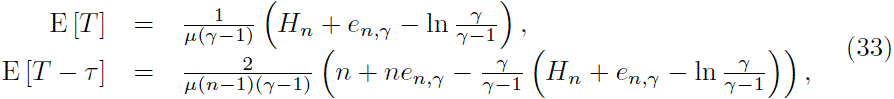

where 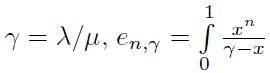 and present the calculated values in Tab. 1. From the formula for the expected quadratic variation,

**Table 1.** Summary of parameters and calculated values for the Hominoid body size analysis.

| Parameter | Phylogeny 1<br>[Purvis, 1995] | Phylogeny 2<br>[Bokma et al., 2012] | DLMH<br>[Bokma et al., 2012] |
| --- | --- | --- | --- |
| $n$ | 5 | 7 | — |
| $\lambda$ | 0.134 | 0.46 | — |
| $\mu$ | 0.037 | 0.43 | — |
| $E[\Upsilon]$ | 3.553 | 20.952 | 3.6 |
| $E[v]$ | 1.448 | 15.639 | — |
| $E[T]$ | 20.831 | 24.935 | 5.25 |
| $E[T - \tau]$ | 12.941 | 19.182 | — |
| $\rho_n$ only gradual change | 0.621 | 0.769 | — |
| $\rho_n$ gradual and punctuated change | 0.615 | 0.766 | — |
| $E[[X]]$ | 1.038 | 1.384 | 0.2845 |
| % anagenetic | 97.11% | 87.21% | 89.31% |
| % cladogenetic | 2.89% | 12.79% | 10.39% |

**Table 2.** Hominoidea body size measurements, read off Fig 2. in [Bokma, 2002].

| Species | body size $[\ln(kg)]$ |
| --- | --- |
| *Gorilla gorilla* | 4.74 |
| *Homo sapiens* | 4.1 |
| *Pongo Pygmaeus* | 4.02 |
| *Pan paniscus* | 3.66 |
| *Pan troglodytes* | 3.62 |

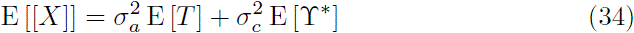

I quantified the amount of change attributed to anagenetic and cladogenetic evolution by

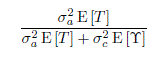

and

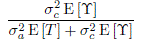

respectively. Table 1 summarizes these values for the studied tree setups.

In his study Bokma [2002] estimated the parameters of anagenetic and cladogenetic change and concluded that one cannot reject the null hypothesis that body size evolution in Hominoidea has been entirely gradual. I will assume here his estimated values of 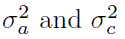 and conditional on them see how punctuated change effects Hominoidea. Therefore some way of working with the number of jumps that occurred is required and there are two possible ways to do this. One is to use the estimated values of E[ϒ] and E[*ν*]. However, since I have a tree available I may try to estimate the number of jumps conditional on the tree for a particular lineage of interest. In this case there is no need to use E[*T*] as the lineage length will be known. I will discuss both approaches and in the second case choose after Bokma et al. [2012] the direct line to modern human (DLMH) as the lineage of interest.

From Tab. 1 One can see that in the case of the first five species phylogeny cladogenetic change decreases the species' similarity by 0.0062 (or by 1%). To get some sort of comparison with the real data (Fig. 2) I considered all 20 pairs of species and from this estimated the correlation as 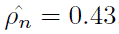. Notice that this is not entirely correct as the sampled pairs are statistically dependent. However, this empirical correlation coefficient is on the same level as the theoretical one of 0.6151. One can also see that the punctuated change consists of 2.9% of the expected change of the phenotype. Therefore even if the estimate of 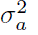 from Bokma [2002] was significant the role of punctuated change would not be that crucial.

Next I studied evolution on the DLMH. Its length is suggested to be between 4.1 and 7.02 Mya [Hobolth et al., 2007, Kumar et al., 2005, Patterson et al., 2006] I chose T ≈ 5.25[Mya] [Bokma et al., 2012, phylogeny in supporting material] to be consistent with the estimated number of speciation events [even though in Purvis, 1995, according to which 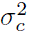 was estimated, 7.04Mya is stated, this however would only increase the contribution of gradual change by about 2.5%].

The estimated expected number of speciation events on the lineage is 3.6 [Bokma et al., 2012] This gives the expected amount of log–body–mass change in the lineage to *Homo sapiens* from our divergence from chimpanzees to be 0.2845 and in Tab. 1 one see that gradual change remains the dominant one but the punctuated change plays some role as well. The two *Pan* species have an average log–body–mass of 3.64ln(*kg*) while *Homo sapiens* has 4.1ln(*kg*). The difference between these two values is 0.46, not very far from 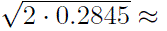 0.754, the square root of the expected total amount of change, occurring on the distance from *Homo sapiens* to the *Pan* clade. We take the square root to have the same unit of measurement, ln(*kg*), in both cases. Actually one would expect the actual difference to be lower than the total change. This is because the real fluctuations are random and will not always be divergent. Sometimes they will be convergent, i.e. the change in both lineages will be sometimes in the same and sometimes in a different direction, therefore some changes will cancel out. The “total change” (or quadratic variation) on the path connecting the two clades on the other hand, will not have this evening out effect, but will first make all change divergent and then sum it.

Comparing the percentage of punctuated change to the case with the random birth–death tree one can see that even though both indicate that punctuated change is not the dominating explanation for evolution, they differ in the estimated magnitude of it. I will compare what happens if instead of the birth–death rates of Purvis et al. [1995] I use λ = 0.46, *μ* = 0.43 and 7 species [after Bokma et al., 2012] In Tab. 1 one can see higher correlation coefficients and the proportion of anagenetic and cladogenetic change is similar to what I obtained from the DLMH calculations. This however should not be that surprising as both the length of the DLMH and the birth and death rates used above were jointly estimated by Bokma et al. [2012].

With this analysis I do not intend to say anything definitive about Hominoid body size evolution. It is only meant to serve as an illustration for the methodology presented here. All of the calculations seem to indicate that evolution in this clade was predominantly driven by gradual change.

## 5 Discussion

Bokma [2002] introduced in his work [but see also Bokma, 2003, 2008, 2010, Mattila and Bokma, 2008, Mooers and Schluter, 1998, Mooers et al., 1999] a modelling approach allowing for both gradual change and punctuated equilibrium. The modelling framework presented here is compatible with his but looks at it from a different perspective. Bokma [2002] was interested in detecting punctuated equilibrium and devised a statistical procedure for it, including confidence intervals. I ask another question: if I know (have estimates of) rates of cladogenetic and anagenetic evolution, can I then quantify (as e.g. fractions/percentages) how much of evolutionary change is due to gradual and how much due to cladogenetic change. These are two different questions. The first is a statistical one, is there enough data to estimate a parameter and how to do it. The second is descriptive, what do parameters mean and what effects do they have on the system under study [but see also Mattila and Bokma, 2008, for quantifying effects of gradual and punctuated evolution].

In Eq. (29) one can see that the magnitude of each type of evolution depends on two components, its instantaneous effect 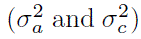 and the time over which each was allowed to act (*T* and ϒ*). If speciation is very rare, then even if 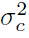 is much larger than 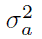, punctuated equilibrium will have a tiny effect on the final evolutionary outcome. However, if the jumps are frequent, cladogenetic evolution can have a very large effect even if its magnitude is small each time it occurs.

I observed in addition that adding a cladogenetic component has the effect of making species less and less similar, as expected from intuition. The magnitude of the cladogenetic effect could give some insights into a trait’s role. As suggested by Mooers et al. [1999] cladogenetic evolution could be linked to traits that are involved in speciation or niche shifts and gradual evolution to traits under continuous selection pressures that often change direction.

I also believe that my characterization of the Ornstein–Uhlenbeck process with jumps is an important contribution to evolutionary modelling. This model is consistent with many theories on trait evolution. It is constrained and so it is well suited to include the fact that the trait has to function with other traits and cannot be arbitrarily large or small as then it would become useless. Of course some freedom to vary around the optimum has to be present, e.g. to allow the organism to adapt if the environment changes. This is exactly how the Ornstein–Uhlenbeck process behaves. At stationarity it is made up of constant, finite variance oscillations around the optimum. If the optimum changes, the trait will be pulled towards the new optimum. If the phenotype changes radically due to e.g. rapid speciational change, it will be pulled back towards the optimum.

Most phylogenetic comparative data sets contain only contemporary measurements. These measurements have accumulated both cladogenetic and anagenetic effects over the whole course of their evolution. Even if the instantaneous effect of cladogenetic evolution is large, if it occurred rarely compared to the time of anagenetic evolution it could be difficult to obtain statistical significance of parameter estimates. Therefore, it is crucial [as also pointed out in Bokma, 2002] to include speciation events resulting in extinct lineages. At each hidden speciation event cladogenetic evolution may have taken place.

From the Hominoid body size example discussed here one can conclude that in the study of punctuated equilibrium if one conditions on the speciation and extinction rates it is important to obtain correct values of them to have the right balance between the time when gradual change took place and the number of opportunities for punctuated change especially with a small number of tip species. I conducted an analysis based on two studies of essentially the same clade but with different birth and death rate estimates and obtained different (though qualitatively similar) proportions. However, one can also see that what enters Eq. (30) is the product 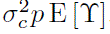. Therefore one is free to vary two components 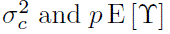 to obtain the cladogenetic contribution. A low jump frequency and a high variance of the jump will give the same effect as a high frequency of low magnitude jumps. Therefore it seems plausible that in a joint estimation procedure even if it would be difficult to reliably estimate the individual parameters 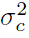, *μ* and λ one still might be able to obtain reasonable values for the contribution of each mode of evolution.

I have presented analytical results for the conditioned Yule model. This model does not allow for extinction but it is a starting point for integrating punctuated equilibrium models of phenotypic evolution with models of tree growth. As I discussed, trees with extinction can easily be handled by simulation methods and a pure birth model can serve as a convenient prior for a Bayesian approach or as the basis for a numerical procedure. In addition to this I have derived the probability generating function for the number of speciation events on a random lineage and for the number of speciation events from the most recent common ancestor of two randomly sampled tip species. These are to the best of my knowledge novel results [but see also e.g. Stadler, 2009, Steel and McKenzie, 2001, for further distributional properties of Yule trees].

## Acknowledgments

I would like to thank Folmer Bokma for his many helpful comments, corrections and reference suggestions that immensely improved the manuscript. I would like to thank Olle Nerman for his suggestions leading to the study of models with jumps at speciation events, and Thomas F. Hansen, Serik Sagitov and Anna Stokowska for many valuable comments and suggestions. I was supported by the Centre for Theoretical Biology at the University of Gothenburg, Stiftelsen for Vetenskaplig Forskning och Utbildning i Matematik (Foundation for Scientific Research and Education in Mathematics), Knut and Alice Wallenbergs travel fund, Paul and Marie Berghaus fund, the Royal Swedish Academy of Sciences, and Wilhelm and Martina Lundgrens research fund. A previous version of this manuscript went into my PhD thesis at the Department of Mathematical Sciences, University of Gothenburg, Gothenburg Sweden.

## A

### Theorem proofs

#### A.1 Proof of Theorem 2.1

##### Proof.

ϒ was written as,

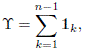

where the indicator random variables **1**_*k*_ represent whether the *k*-th coalescent event is on the sampled lineage and by properties of the Yule tree they are independent and equal one with probability 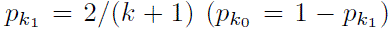. Therefore,

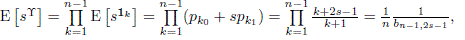

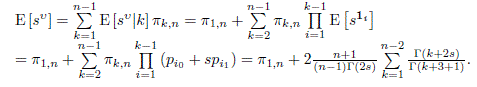

I assumed *n* > 2 (if *n* = 2 then by definition *ν* = 0 and so for all 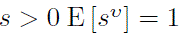) and I use the property [see also Bartoszek and Sagitov, 2012],

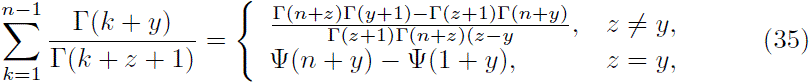

where Ψ(*z*) is the polygamma function defined as 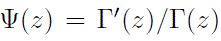. In the case of *z* ≠ *y* the formula above is verifiable by induction and if *z* = *y* the formula can be either calculated directly from the left side or as the limit of the right side *z* → *y*.

This property and that 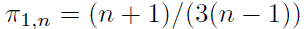 gives when *s* ≠ 1.5,

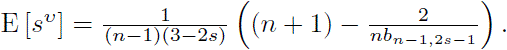

When *s* = 1.5 I get,

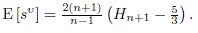

#### A.2 Proof of Theorem 3.2

##### Proof.

Let 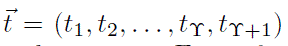 be the between speciation times on a randomly chosen lineage, see Fig. 8 for illustration. For mathematical convenience I set the speciation rate λ = 1 as changing λ is equivalent to rescaling the branch lengths by its inverse and does not effect the topology (ϒ and *ν* here) because I have conditioned on *n*. Therefore a Yule–Ornstein–Uhlenbeck–jumps model for the extant species trait sample with parameters 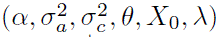 is equivalent to one with parameters 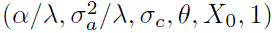. By 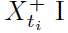 I will denote the value of the process just after (+ indicating I include the jump if it occurred) the node ending the branch corresponding to duration *t*_*i*+1_, notice that 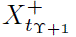 will be the value just after an extent species and 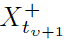 will be the value just after the node being the most recent common ancestor of two randomly sampled species. If I don't include the +, i.e. *X_t_i__* I mean the value of the process at the respective node. As in the proof of Theorem 2.1 1*k* is the 0-1 random variable indicating whether the *k*-th (*k* = 1,…, *n* – 1) coalescent event is on the sampled lineage, equalling 1 with probability 2/(*k* + 1). The expectation of an extant species is,

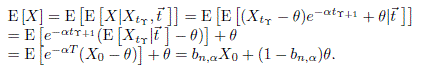

I now turn to the variance,

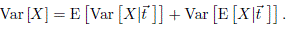

I consider the most complicated term,

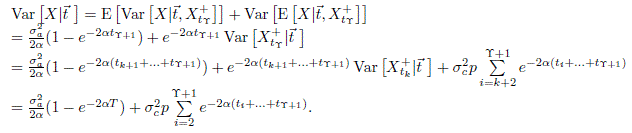

I will use the equality, for *y* > 0, obtainable from Eq. (35),

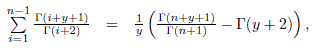

to calculate,

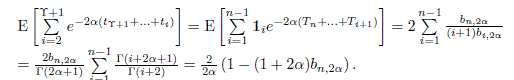

Using the above and

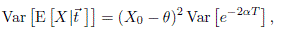

I arrive at,

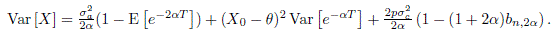

Writing 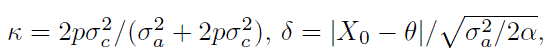

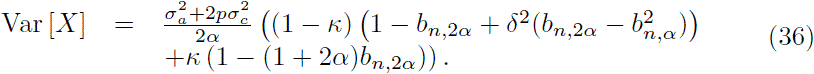

In the same manner I will calculate the covariance between two randomly sampled tip species (with 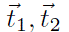 being the vectors of between speciation times on the respective species’ lineage),

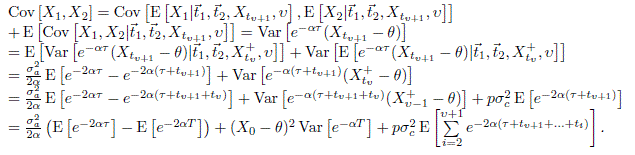

Let *κ* denote number of the speciation event when the two randomly sampled tips coalesced and I consider for *y* > 0,

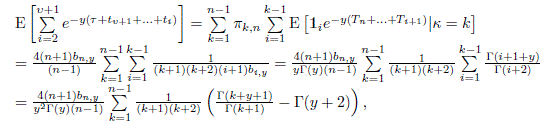

equalling by Eq. (35) for 0 < *y* ≠ 1

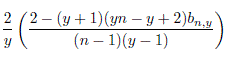

and for *y* = 1 this is,

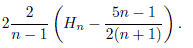

Putting all of the above together I obtain,

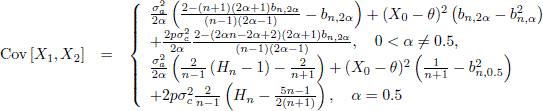

and with *κ* and *δ*,

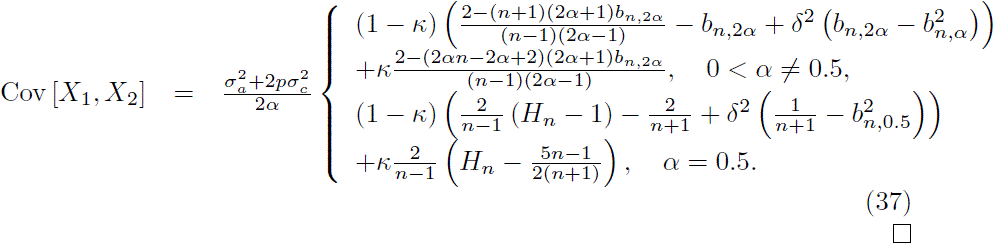

### B Quadratic variation

In order to quantify how much a function (for example a trait as a function of time evolving on a lineage) changed over time one needs to have some measure of change. It is not sufficient to compare the current function (trait) value (or average of trait values) with that of the value of the function at the origin (ancestral trait). It can happen that the function might have over time made large excursions but just by chance is currently at a value close to the origin, especially if the expectation of the (random) function equals that of the original value, like in the Brownian motion (neutral evolution) one.

One therefore needs some measure that will consider not only how the function compares to what it was at some time in the past but will also (or rather primarily) take into account how diverse its path was. One such possible measure is the quadratic variation of a function.

#### Definition.

*The quadratic variation of a function f*(*t*) *is defined as,*

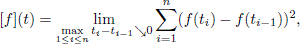

*where* 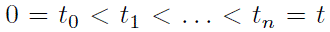 *are finer and finer partitions of the interval* [0,*t*].

The advantage of using quadratic variation to quantify the phenotype’s change is that it is mathematically easy to work with and will be consistent with the ideas and results of Bokma [2002], Mattila and Bokma [2008]. The quadratic variation is a property of a function and so can be applied to a trajectory of a stochastic process. In fact one can show [see e.g. Klebaner, 2007] that the quadratic variation of a stochastic process defined by an SDE of the form,

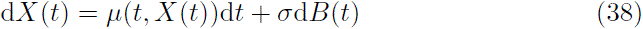

where *B*(*t*) is a standard Brownian motion and with appropriate conditions on *μ*(·,·) will be 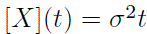. This is rather remarkable as the quadratic variation is a property of the process’ trajectory (a random function) but it turns out to be a deterministic function. From the definition one can see that the quadratic variation takes into account all the local fluctuations of the phenotype and then sums them up to get the accumulated fluctuation. The considered above family of SDEs is sufficient for my purposes as it encompasses the two most widely used evolutionary models, Brownian motion, *μ* ≡ 0, and Ornstein–Uhlenbeck process, 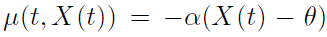. For further properties of the quadratic variation the reader is referred to stochastic calculus literature [e.g. Klebaner, 2007, Medvegyev, 2007, Øxendal, 2007].

